# Functional Redundancy of the Posterior Hippocampi, but not Anterior Hippocampi or Left Frontal Cortex, is Disrupted in Pathological Brain Aging

**DOI:** 10.1101/2022.06.18.496543

**Authors:** Jenna K. Blujus, Michael W. Cole, Elena K. Festa, Stephen L. Buka, Stephen P. Salloway, William C. Heindel, Hwamee Oh, the Alzheimer’s Disease Neuroimaging Initiative

## Abstract

As prevalence rates of Alzheimer’s disease (AD), the leading cause of dementia, are projected to more than double by 2050, emphasis has been placed on early intervention strategies that target resilience mechanisms to delay or prevent the onset of clinical symptoms. Several neural mechanisms underlying brain resilience to AD have been proposed, including redundant neural connections between the posterior hippocampi (HC) and all other brain regions, and global functional connectivity of the left frontal cortex (LFC). It remains unknown, however, if regional redundancy of the HC and LFC underscores neural resilience in the presence of AD pathologies. From the ADNI database, 363 cognitively normal older adults (CN) (N = 220; 36% A*β*+) and patients with Mild Cognitive Impairment (MCI) (N = 143; 51% A*β*+) were utilized. Regional redundancy was calculated from resting state fMRI data using a graph theoretical approach by summing the direct and indirect paths (path lengths=1-4) between each ROI and its 262 functional connections. The results showed that A*β*-status significantly disrupted posterior HC, but not anterior HC or LFC, redundancy. A*β*- groups showed higher redundancy of the bilateral posterior HC than A*β*+. In regard to redundancy-cognition relationships, higher posterior HC redundancy was related to better episodic memory performance, an effect which was primarily driven by the A*β*- group. Despite the positive relationship between posterior HC redundancy and cognition, we did not find compelling evidence that redundancy of the posterior HC serves in a resilience manner, as posterior HC redundancy did not moderate the potentially deleterious relationship between A*β* deposition and cognition. No relationships were found between anterior HC or LFC redundancy and cognitive performance. Together, these findings suggest that redundancy of the LFC does not underpin its role in resilience and that posterior HC redundancy may capture disruptions to network connectivity that occur as a result of A*β* deposition.

## Introduction

Alzheimer’s disease (AD), the leading cause of dementia, is pathologically characterized by *β*-amyloid (A*β*) plaques and neurofibrillary tangles, accompanied by progressive cognitive decline (Braak & Braak, 1991; Querfurth & LaFerla, 2010). Neuropathological and *in vivo* neuroimaging evidence has demonstrated that around 30% of clinically normal adults exhibit significant loads of AD neuropathology in the brain (Esiri et al., 2001; Jagust, 2013; Morris et al., 1996; Price & Morris, 1999), representing a stage now called preclinical AD (Jack Jr et al., 2018; Sperling, Karlawish, & Johnson, 2013). Such findings of a disconnection between neuropathology and clinical status led to the hypotheses collectively considered as reserve and resilience (Arenaza-Urquijo & Vemuri, 2018; Katzman et al., 1988; Stern, 2002, 2009). Reserve and resilience hypotheses postulate that certain neural mechanisms, which are potentially multiple yet complementary, exist to enhance an individual’s ability to cope with developing pathology (Katzman et al., 1988; Stern, 2002, 2009; Stern, Barnes, Grady, Jones, & Raz, 2019). These neural mechanisms supposedly underlying reserve and resilience in the presence of pathology are further thought to be shaped by enriching experiences accumulated across the lifespan, such as higher education, literacy, physical exercise and social engagement (Cabeza et al., 2018; Stern, 2002). Given that no effective treatment for AD is readily available, efforts on prevention and early intervention targeting neuroprotective or resilience mechanisms have been emphasized to delay or prevent a large proportion of dementia cases (Livingston et al., 2020).

Several neural mechanisms underlying resilience have been proposed. One is an increased neural activity measured by positron emission tomograpy (e.g., [18F]Fluorodeoxyglucose PET), regional cerebral blood flow, or functional magnetic resonance imaging (fMRI) during tasks. The increased neural activity, commonly construed as neural compensation, has been seen in particular in the frontoparietal cortex, which is accompanied by temporoparietal hypometabolism (Stern, Alexander, Prohovnik, & Mayeux, 1992) or hypermetabolism despite the presence of brain A*β* pathology (Benzinger et al., 2013; Johnson et al., 2014; Oh, Habeck, Madison, & Jagust, 2014; Oh, Madison, Baker, Rabinovici, & Jagust, 2016; Ossenkoppele et al., 2014; Stern et al., 1992). Specifically, a higher level of education is associated with decreased neural activity in the temporoparietal cortices and increased neural activity in frontoparietal cortex (Oh, Razlighi, & Stern, 2018; Stern et al., 1992). Such increased neural activity has been accompanied by either better memory performance or nondemented cognitive status (Ossenkoppele et al., 2014). Based on these observations, it was hypothesized that the brain regions with an increased neural activity may play a role of compensating for the disrupted functions in the temporoparietal cortices and their connected brain regions such as hippocampus.

Network-based mechanisms, measured using resting state fMRI, have also been proposed to underlie resilience in brain aging, including global connectivity of the left frontal cortex (LFC) (Franzmeier et al., 2017, 2018) and functional redundancy of the posterior hippocampi (Langella, Sadiq, Mucha, Giovanello, & Dayan, 2021). The LFC is a hub region involved in cognitive control and acts to flexibly and adaptively recruit additional neural networks to regulate behavior (Cole, Yarkoni, Repovš, Anticevic, & Braver, 2012; Cole et al., 2013). Global connectivity of the LFC was positively associated with education and was shown to mitigate the slope of memory decline in patients with Mild Cognitive Impairment (MCI) and delay cognitive impairment in autosomal dominant and sporadic Alzheimer’s disease (Franzmeier et al., 2017, 2018). In addition to LFC connectivity, a measure of hippocampal redundancy, capturing the topography of functional connections to the hippocampus (HC), was recently proposed as a resilience mechanism in healthy aging and a prodromal stage of AD (Langella et al., 2021). Redundancy indicates that there are multiple parallel pathways that exist to functionally connect any two brain regions in a network, thereby providing a potential fail-safe mechanism within the system in the event of failure or dissolution of a functional connection between brain two brain regions. In the context of AD, alternative connections between brain regions may serve to prevent total functional loss in the presence of developing neuropathology or neurodegeneration. Recent work demonstrated that redundancy of the left and right posterior hippocampus (HC), but not the anterior HC, was higher in cognitively normal (CN) than patients with MCI (Langella et al., 2021). Further, higher posterior HC redundancy was associated with better episodic memory performance, a relationship which was seen in CN and early MCI. These findings provided preliminary insight that redundancy, particularly of the posterior HC, may confer resilience in prodromal stages of AD (Langella et al., 2021).

Though both HC redundancy and global LFC connectivity are compelling correlates of cognitive resilience in normal and pathological brain aging, a few elements remain to be clarified. First, in regard to HC redundancy, it remains to be tested whether anterior or posterior HC redundancy is affected by AD neuropathology. Further, it is not clear if HC redundancy plays a resilience role at the earliest stages of AD pathologies through moderating the potential deleterious effect of AD neuropathology on cognitive performance. In regard to LFC, the current literature suggests that the mechanism by which LFC connectivity confers cognitive resilience is through the strength of its immediate functional connections dispersed throughout the brain (Cole et al., 2012; Franzmeier et al., 2017). What remains unclear, however, is whether the neuroprotective effects of the LFC are also mediated by additional mechanisms, such as functional redundancy. Though redundancy and global connectivity are complimentary metrics, in that they both capture connectivity of the LFC, these metrics measure qualitatively different aspects of connectivity. For example, global connectivity measures the strength of the connections between the LFC and its direct functional neighbors, while redundancy encapsulates the global topology of LFC connectivity, representing both the direct and indirect functional connections. Thus, further investigation is warranted to discern if redundant, non-direct functional connections of the LFC also underlie its role in resilience.

The purpose of the current study was to determine if regional redundancy (i.e., HC and LFC) underscores resilience in the presence of AD pathologies. Using resting state fMRI data from the Alzheimer’s Disease Neuroimaging Initiative (ADNI) database, redundancy was estimated using a graph theoretical approach in five regions of interest (i.e., left and right anterior HC, left and right posterior HC, LFC) in CN and MCI groups characterized by A*β* positivity status. Based on previous work, we expected that posterior HC, but not anterior HC or LFC, would be lower in MCI than CN (Franzmeier et al., 2018; Langella et al., 2021). Further, we expected that if regional redundancy plays a role in resilience in the earliest stages of developing AD pathologies, redundancy levels in the CN A*β*+ groups would be greater than MCI, regardless of A*β*-status, and exhibit comparable or higher levels than that of the CN A*β*- group. It was also hypothesized that if regional redundancy serves in a resiliency manner, the potential negative relationship between cognition and pathology would be mitigated by higher redundancy. If HC redundancy reflects healthy brain aging status, however, higher HC redundancy would be observed in A*β*- than A*β*+ groups.

## Method

### Participants

The data utilized for analysis were obtained from the Alzheimer’s Disease Neuroimaging Initiative (ADNI) database. Launched in 2003 as a public-private partnership, ADNI is led by Principal Investigator Michael W. Weiner, MD. A goal of ADNI has been to test whether serial magnetic resonance imaging (MRI), positron emission tomography (PET), other biological markers, and clinical and neuropsychological assessment can be combined to measure the progression of MCI and early AD. For more information, see www.adni-info.org.

A total of 370 CN and MCI participants were selected for analysis from the ADNI database. The inclusion criteria for each diagnostic group were determined by ADNI. CN participants presented with no memory complaints, had MMSE performance between 24-30, Clinical Dementia Rating (CDR) performance of 0, and did not meet diagnostic criteria for MCI or AD. MCI participants presented with a subjective memory concern, MMSE performance between 24-30, CDR score of 0.5, preserved functioning, and did not meet criteria for diagnosis of AD. Participants selected for analysis additionally underwent AV45 PET within one year of the resting state fMRI session. A*β*-status was determined using a threshold of 1.11 Standardized Uptake Value Ratios (SUVR) (normalized to the cerebellum), which is recommended by ADNI for cross-sectional studies (Jones et al., 2016). Individuals above the A*β* criterion were deemed A*β*-positive (A*β*+) while those below were deemed A*β*-negative (A*β*-). Seven participants were missing *APOE4* genotype information and thus were excluded from analysis, resulting in a total sample of 363 individuals. See Table 1 for sample demographics.

**Table 1.** Demographic Characteristics.

| | CN Aβ- | CN Aβ+ | MCI Aβ- | MCI Aβ+ |
| --- | --- | --- | --- | --- |
| N | 140 | 80 | 70 | 73 |
| Age Range | 55.8-91.3 | 61.1-91.5 | 59.7-91.6 | 56.7-93.2 |
| Age<sup>a</sup> | 72.91 (6.94) | 75.77 (6.92) | 75.49 (7.99) | 74.54 (7.70) |
| Sex (% female) | 55.00 | 56.25 | 42.86 | 47.95 |
| Education (years) | 16.73 (2.33) | 16.52 (2.59) | 16.17 (3.08) | 16.10 (2.55) |
| *APOE*ε4 (% carriers)<sup>c</sup> | 20.71 | 47.50 | 17.14 | 61.64 |
| MMSE<sup>a</sup> | 29.11 (1.11) | 28.78 (1.43) | 28.37 (1.69) | 27.33 (2.04) |
| Memory (composite)<sup>a</sup> | 1.05 (0.62) | 0.94 (0.62) | 0.60 (0.67) | 0.10 (0.63) |
| EF (composite)<sup>b</sup> | 1.11 (0.77) | 0.94 (0.79) | 0.42 (0.81) | 0.29 (0.95) |
| FD | 0.18 (0.09) | 0.20 (0.20) | 0.20 (0.10) | 0.20 (0.11) |

Note. Values presented as $M \pm SD$ unless otherwise specified. MMSE = Mini Mental State Exam; EF = executive function; FD = framewise displacement.
<sup>a</sup> Significant interaction effect between diagnosis and Aβ-status, $p$ 's < .05.
<sup>b</sup> Significant main effect of diagnosis, $p$ 's < .05.
<sup>c</sup> Significant main effect of Aβ-status, $p$ 's < .05.

### Data Acquisition and Preprocessing

Resting state fMRI data from ADNI phases 2 and 3 (basic sequence) were used in the current analysis. The MR acquisition protocols employed across phases were common to all sites and adapted for each scanner (volumes=140-200, TR=3000ms, TE=30ms, flip angle=80-90°, slice thickness=3.3-3.4mm).

Resting state fMRI data were preprocessed using fMRIPrep 20.2.6 (Esteban et al., 2019), which is based on Nipype 1.7.0 (Gorgolewski et al., 2011). The functional data were slice-time corrected (*3dTshift*; AFNI v16.2.07) (Cox & Hyde, 1997), motion corrected (*mcflirt* FSL v5.0.9) (Jenkinson, Bannister, Brady, & Smith, 2002), and co-registered to the anatomical T1-weighted scan using boundary based registration with six degrees of freedom (*bbregister* ; Freesurfer v6.0.1) (Dale, Fischl, & Sereno, 1999; Greve & Fischl, 2009). Several confounding time-series were calculated based on the preprocessed fMRI blood oxygenation level dependent (BOLD) signals: framewise displacement (FD) (Power et al., 2014) and anatomical component-based regressors computed separately for CSF and WM (aCompCor) (Behzadi, Restom, Liau, & Liu, 2007). Volumes that exceeded a threshold of 0.5 mm FD were noted as motion outliers (Power, Barnes, Snyder, Schlaggar, & Petersen, 2012; Power et al., 2014). All transformations (i.e. head-motion transform matrices and co-registrations to anatomical and template MNI spaces) were performed with a single interpolation using *antsApplyTransforms* (ANTs v2.3.3) (Avants, Epstein, Grossman, & Gee, 2008), configured with Lanczos interpolation (Lanczos, 1964).

Following preprocessing, AFNI v21.2.04 (Cox & Hyde, 1997) was used for denoising. Voxel timeseries were scaled to a mean of 100 (*3dTstat, 3dcalc*). Nuisance regression included volumes identified as non-steady state outliers by fMRIprep, six realignment parameters and their first temporal derivatives, and the first five anatomical principal components from WM and CSF (Ciric et al., 2017; Matijevic, Andrews-Hanna, Wank, Ryan, & Grilli, 2021). The data were linearly detrended and a bandpass filter (0.008 - 0.09 Hz) was applied.

Groups were matched in terms of average motion (FD). There were no diagnostic differences in FD, *F* = 0.47, *p* = .496, no differences in FD due to A*β*-status, *F* = 0.57, *p* = .453, and no significant interaction of diagnosis and A*β*-status on FD, *F* = 0.87, *p* = .350.

### Brain Network Construction

Average timeseries were extracted from 263 functionally-defined ROIs (Seitzman et al., 2020), which provided thorough coverage of cortical, subcortical, and cerebellar regions (Figure 1A). The timeseries of each ROI pair were correlated and Fisher z-transformed to form a 263 x 263 ROI functional connectivity matrix (i.e., graph) in each participant. For redundancy calculations, matrices were binarized using a proportional threshold technique and self-connections were set to zero. To ensure that differences in redundancy were not dependent on graph density (Rubinov & Sporns, 2010), graphs were thresholded across a wide range (2.5-25%, steps of 2.5%) (Langella et al., 2021) and redundancy was calculated at each threshold. To summarize redundancy properties across the wide range of densities sampled in the current analysis (2.5-25%), area under the curve (AUC) was calculated to represent redundancy as a single scalar metric (K. Li et al., 2015). AUC has widely been applied in analyses of graph-based networks and is sensitive to detect pathological alterations in network topology (K. Li et al., 2015; W. Li et al., 2020; Luo et al., 2015; Zhang et al., 2011).

**Figure 1.**
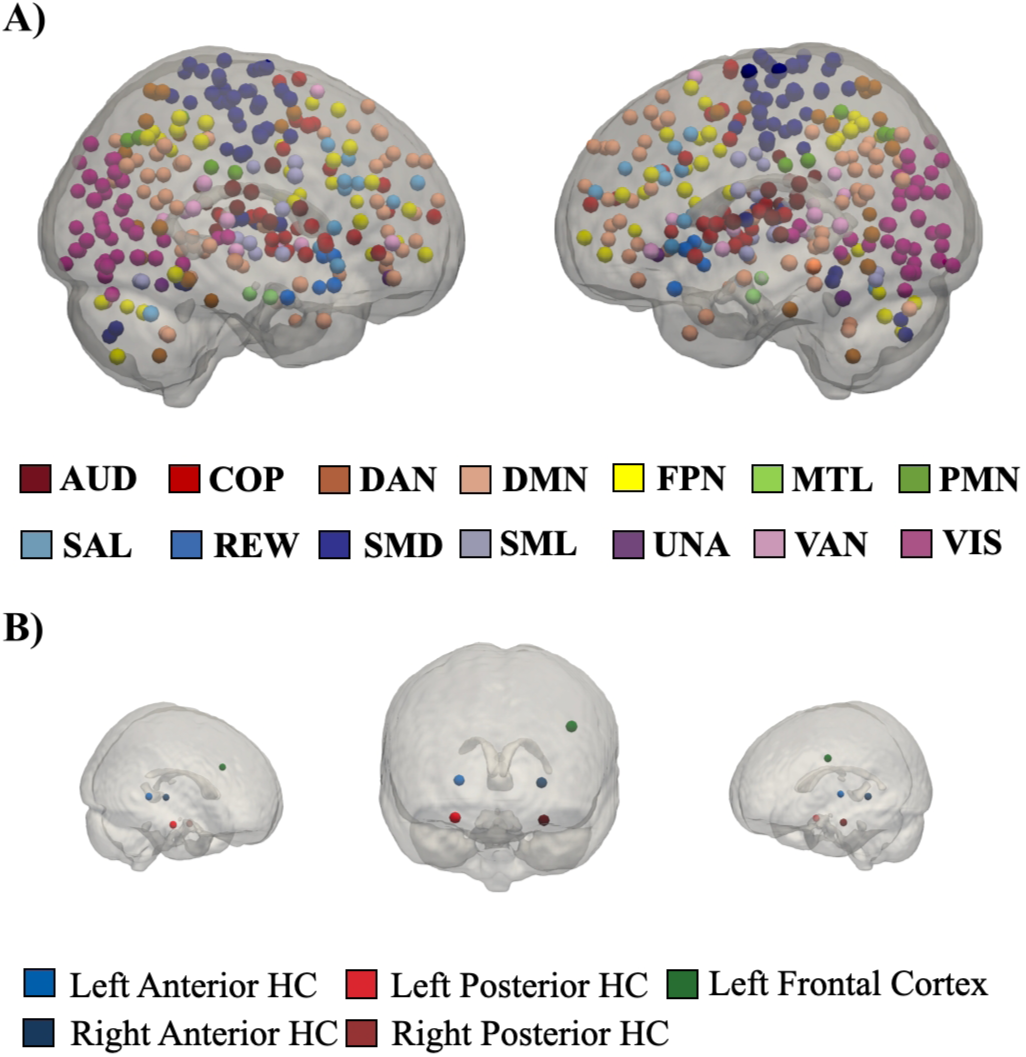
Brain nodes and a priori defined ROIs. (A) 263 nodes distributed across 13 networks displayed in glass brain. (B) A priori defined ROIs displayed in radiological view. ROIs included the anterior and posterior bilateral hippocampi and the left frontal cortex. HC = hippocampus; AUD = auditory network; COP = cingulo opercular network; DAN = dorsal attention network; DMN = default mode network; FPN = frontoparietal network; MTL = medial temporal lobe network; PMN = parietomedial network; SAL = salience network; REW = reward network; SMD = somatomotor network dorsal; SML = somatomotor network lateral; UNA = unassigned; VAN = ventral attention network; VIS = visual network.

### Redundancy

Global redundancy *(R_global_)* captures the total number of repetitive (e.g., direct and indirect) connections between node pairs. A 263 x 263 redundancy matrix was formed by summing the total paths of lengths (*l*)=1 to L for each node pair (Di Lanzo, Marzetti, Zappasodi, De Vico Fallani, & Pizzella, 2012; Langella et al., 2021):

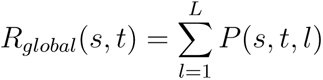

L (maximum path length) was set to four based on computational demands and guidelines of previous work (Di Lanzo et al., 2012; Langella et al., 2021). Higher R_global_ of node pair (*s*, *t*) suggests that there are many alternative pathways connecting nodes *s* and *t*.

To calculate the R_global_ of each region, row averages of the redundancy matrix were calculated. The resulting 263 x 1 column vector of redundancy values represents the average number of total connections (e.g., direct and indirect) between each ROI and its 262 connections. Five regions were defined *a priori* and selected for analysis including the left and right posterior and anterior HC (Langella et al., 2021) and the LFC (Franzmeier et al., 2018) (MNI, x=-41.06, y=5.81, z=32.72) (Figure 1B).

### Direct and Indirect Paths

To further discern whether diagnostic and/or A*β*-status differences in R_global_ were attributed to direct or indirect connections, redundancy components (i.e., direct vs. indirect) were calculated separately. The total number of direct connections between node pairs (R_direct_) was quantified using node degree or the sum of each node’s immediate connections (*a; path length = 1*) (Rubinov & Sporns, 2010):

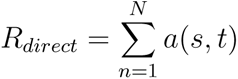

R_indirect_ represented the sum total of all indirect paths connecting node pairs (path length=2-4). Thus, the R_indirect_ matrices excluded direct connections between each ROI-node pair.

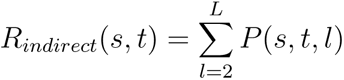

To calculate the R_indirect_ of each region, row averages of the R_indirect_ matrix were calculated. The resulting 263 x 1 column vector of R_indirect_ values represents the average number of total indirect connections between each ROI and its 262 connections. R_indirect_ values from the five *a priori* defined regions were carried forward to analysis.

### Cognitive Performance

To explore redundancy-cognition relationships and the potential role of redundancy as a resilience factor, two domains of cognition affected early in AD were examined, including episodic memory and executive functioning performance. Composite measures of memory and executive functioning were computed by ADNI investigators using the item response theory framework (Gibbons et al., 2012). The memory composite was formed from measures including the RAVLT (learning trials 1-5, interference, immediate recall, delayed recall, recognition), ADAS-Cog (trials 1-3, recall, recognition present, recognition absent), Logical Memory (immediate, delayed), and MMSE (recall of ball, flag, tree word items). The executive function composite included performance from clock drawing, Trails (A, B), Category Fluency (animals, vegetables), WAIS-R Digit (symbol), and Digit Span (backwards). The composite variables were standardized to have a mean of 0 and standard deviation of 1 with higher scores representing better performance.

A high proportion (99%) of the current sample completed a cognitive assessment within six months of their MR scans. One participant was missing data for memory testing and two participants were missing data for executive function testing and thus, were excluded separately for each cognitive domain analysis. The final sample consisted of 362 to explore memory-redundancy relationships and 361 to explore executive function-redundancy relationships.

### Statistical Analysis

Statistical analyses were conducted using R (4.1.0). Demographic characteristics were compared between groups using the *aov_car* function from the *afex* package. Chi-square tests (*chisq.test*) were utilized to examine the influence of diagnosis and A*β*-status on sex and *APOE4* carrier distribution.

To examine the effect of diagnosis and A*β*-status on regional redundancy metrics (i.e., R_global_, R_direct_, R_indirect_ summarized by AUC), non-parametric ANCOVA models with 10,000 permutations were used for each ROI. Age, sex, education, and *APOE4* genotype were included in each model as covariates.

#### Moderation Analysis

Moderation analyses were employed to investigate whether the relationship between the degree of A*β* deposition and regional redundancy was dependent on diagnostic group or A*β*-status. Multiple linear regression models were explored using the *lm* function with ROI redundancy serving as the dependent variable and A*β* SUVR as the predictor, covarying for age, sex, education, and *APOE4* genotype. To examine the moderating effect of diagnostic group, an interaction term of A*β* SUVR and diagnosis was entered into the model. To examine the moderating effect of A*β*-status, separate models were conducted in each diagnostic group (CN, MCI) and included an interaction term of A*β* SUVR and A*β*-status. Conditional effects of the moderation effects were explored using the *probe_interaction* function. Continuous variables were standardized prior to the regression, such that each variable had a mean of 0 and standard deviation of 1. Regression terms were corrected for multiple testing using FDR.

#### Redundancy Relationships with Education

Previous work provided preliminary evidence that posterior hippocampal redundancy may underlie resilience in brain aging (Langella et al., 2021). To further probe this relationship, we examined the face validity of redundancy as a resilience measure by examining its relationship with a proxy of cognitive resilience, namely years of education, for each *a priori* defined ROI. Specifically, multiple linear regression models were formed for each ROI, with years of education serving as the dependent variable and ROI redundancy as the predictor. Each model covaried for age, sex, and *APOE4* carrier status. Continuous variables were standardized prior to the regression.

#### Redundancy Cognition Relationships

To examine the relationship between regional redundancy and cognition, multiple linear regression models were formed for each domain (memory, executive function), with ROI redundancy as the predictor. Each model covaried for age, sex, and *APOE4* carrier status. Continuous variables were standardized prior to the regression.

#### Test of Cognitive Benefit Criterion

The cognitive benefit criterion of resilience posits that neural substrates of resilience will be related to better cognitive performance in the face of brain pathology (Franzmeier et al., 2017). To test this, we employed multiple linear regressions to examine the moderating effect of redundancy on the relationship between the degree of A*β* deposition and cognitive performance (episodic memory, executive functioning). Tests were run separately for each cognitive domain (executive function, memory) and diagnostic group (CN, MCI). For each model, the cognitive composite score served as the dependent variable and A*β* SUVR and redundancy served as the predictor. To examine the moderating effect of redundancy, for each ROI, a redundancy-SUVR interaction term was added to the model. Age, sex, and *APOE4* carrier status served as covariates.

## Results

### Participant Characteristics

See Table 1 for a report of demographic variables by diagnostic group and A*β*-status. The main effects of diagnosis, *F* = 0.71, *p* = .400, and A*β* status, *F* = 1.44, *p* = .232, on age were not significant. However, there was a significant interaction between diagnosis and A*β*-status on age, *F* = 5.70, *p* = .018. Post-hoc tests revealed that CN A*β*- were younger than MCI A*β*-, *p* = .017, and CN A*β*+, *p* = .006. There was a trend-level effect of diagnosis on years of education, *F* = 3.05, *p* = .082, such that CN had higher years of education than MCI. There were no significant differences in years of education by diagnostic group, *F* = 0.24, *p* = .621, and no interaction between diagnosis and A*β*-status, *F* = 0.05, *p* = .821. There was a trend-level effect of diagnosis on sex distribution, *χ*^2^(1) = 3.08, *p* = .079, such that CN had a higher percentage of females in the sample than MCI. A*β* groups did not differ in sex distribution, *χ*^2^(1) = 0.02, *p* = .885. There was a trend-level relationship between diagnosis and *APOE4* carrier distribution, *χ*^2^(1) = 3.00, *p* = .083, such that MCI had a higher percentage of *APOE4* carriers than CN. Further, A*β*+ had a higher percentage of *APOE4* carriers than A*β*-, *χ*^2^(1) = 45.92, *p <* 001. For MMSE performance, both the main effects of diagnosis, *F* = 42.88, *p <* .001, and A*β*-status, *F* = 16.63, *p <* .001, were significant, as well as the interaction diagnosis and A*β*-status on performance, *F* = 4.52, *p* = .034. Regardless of A*β*-status, CN had better MMSE performance than MCI, *p*’s *≤* .001. Within MCI, A*β*- had better MMSE performance than A*β*+, *p <* .001

### Diagnostic and Amyloid Group Differences in Cognitive Performance

See Table 1 for mean cognitive performance by diagnostic group and A*β*-status. In terms of memory performance, both the main effects of diagnosis, *F* = 87.00, *p <* .001, and A*β*-status, *F* = 18.94, *p <* .001, were significant, as well as the interaction of diagnosis and A*β*-status on performance, *F* = 8.11, *p* = .022 (Figure 2A). Within CN, there were memory performance did not differ by amyloid-status, *p* = .243. Though within MCI, A*β*- had better memory performance than A*β*+, *p <* .001. For executive function, there was a significant main effect of diagnosis, such that CN had better executive functioning than MCI, *F* = 55.64, *p <* .001 (Figure 2B). Further, there was a trend-level main effect of A*β*-status on executive functioning performance, *F* = 3.02, *p* = .083, where A*β*- performed better than A*β*+. The interaction between diagnosis and A*β*-status on executive function performance was not significant, *F* = 0.03, *p* = .853.

**Figure 2.**
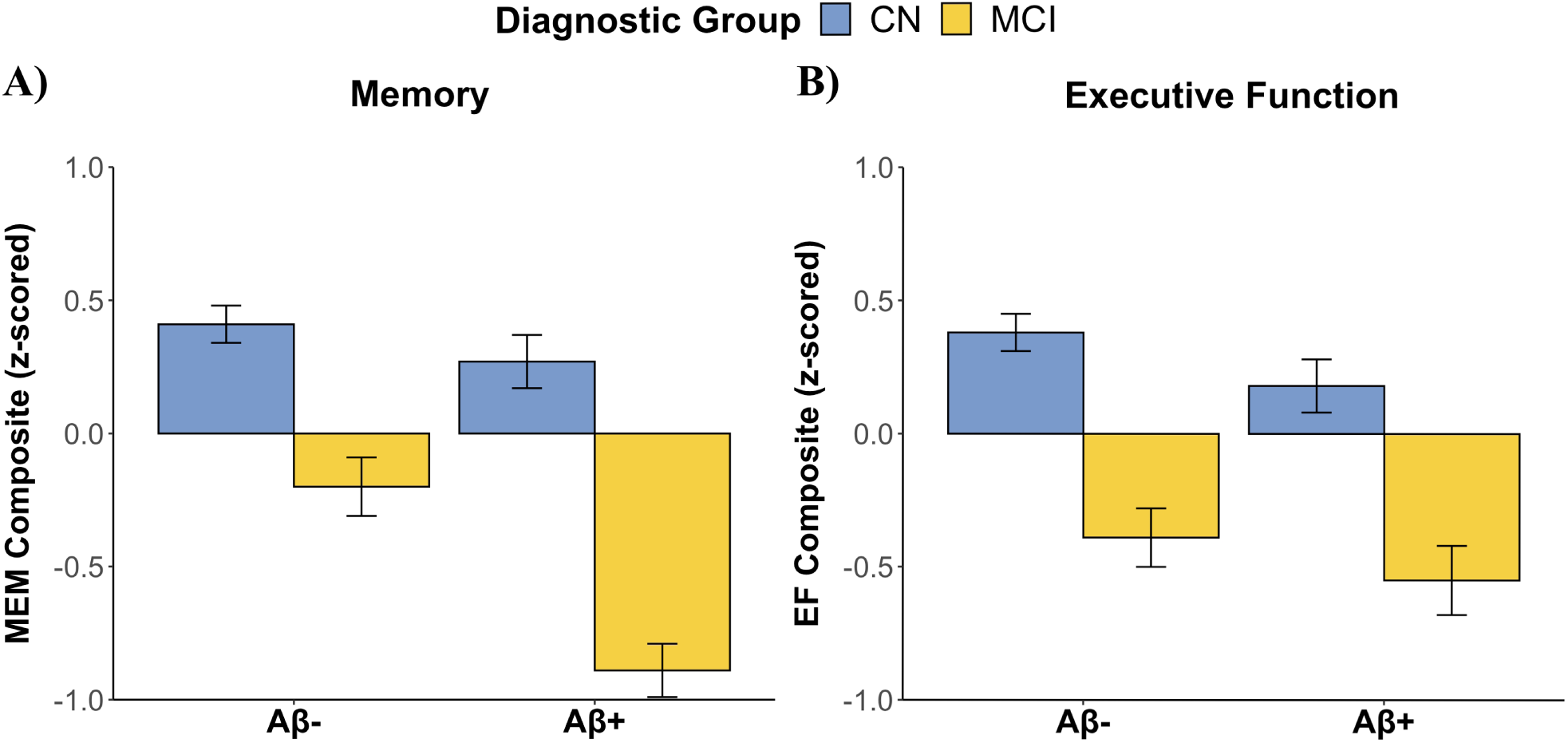
Cognitive performance by group. (A) There was an interaction between diagnosis and Aβ-status on memory performance such that MCI Aβ- had higher memory performance than MCI Aβ+. (B) There were diagnostic differences in executive function performance such that CN had better executive function performance than MCI. There was a trending effect of Aβ-status on executive function performance where Aβ- had better executive function than Aβ+.

### Amyloid-Status Influences Posterior HC Redundancy

We examined the effects of diagnosis and A*β*-status on redundancy (i.e., R_global_) in five *a priori* defined regions of interest, including bilateral posterior hippocampi, bilateral anterior hippocampi, and left frontal cortex. For the posterior hippocampi, there were no significant diagnostic differences in R_global_ (*F* = 0.20, *p* = .659 for left posterior HC; *F* = 0.003, *p* = .954 for the right posterior HC). However, R_global_ differed by A*β*-status in the left posterior HC, *F* = 4.64, *p* = .032, and the right posterior HC, *F* = 6.15, *p* = .014, such that A*β*- had higher posterior HC redundancy than A*β*+ (Figure 3C,D). There were no significant interactions between diagnosis and A*β*-status on R_global_ for the left posterior HC, *F* = 2.25, *p* = .136, or right posterior HC, *F <* 0.01, *p* = .995.

**Figure 3.**
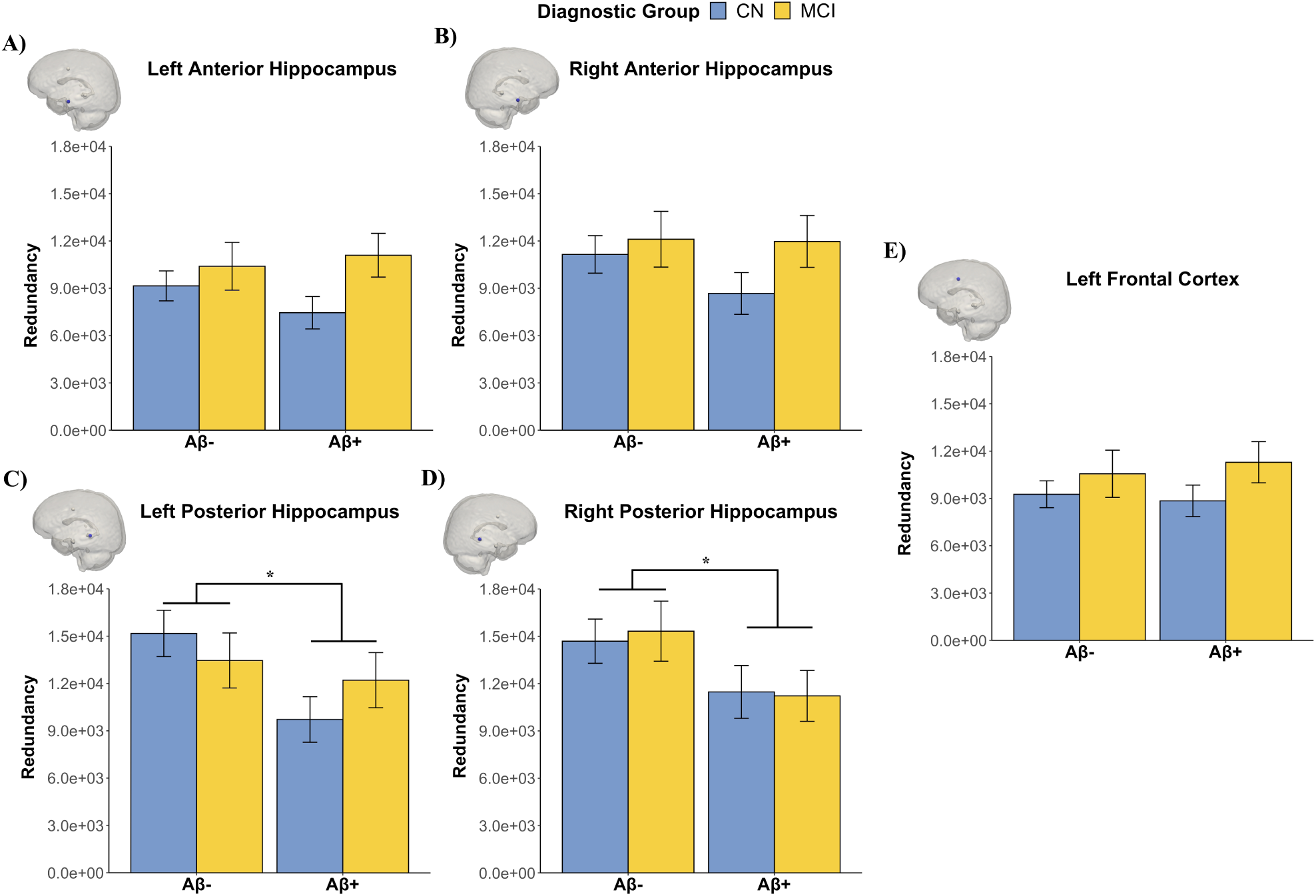
Posterior HC redundancy differs by Aβ-status. Plotted are the diagnostic group by Aβ-status differences in redundancy of the (A) left anterior HC, (B) right anterior HC, (C) left posterior HC, (D) right posterior HC, and (E) left prefrontal cortex. Aβ-status effects were evident in the bilateral posterior HC such that Aβ- had higher redundancy than Aβ+. *p < .05

For the anterior hippocampi, there were trend-level diagnostic differences in left anterior HC R_global_, *F* = 3.61, *p* = .055, such that MCI had higher R_global_ than CN (Figure 3A). There were no diagnostic differences in right anterior HC R_global_, *F* = 2.04, *p* = .153. Further, neither the right or left anterior HC showed differences in R_global_ by A*β*-status (*F* = 0.13, *p* = .712 for left anterior HC; *F* = 1.22, *p* = .269 for right anterior HC, Figure 3B) nor interactions of diagnosis and A*β*-status on R_global_ (*F* = 1.09, *p* = .300 for left anterior HC; *F* = 0.81, *p* = .374 for right anterior HC).

In regard to the LFC (Figure 3E), there were neither main effects of diagnosis, *F* = 1.74, *p* = .184, or A*β*-status, *F* = 0.0003, *p* = .987, nor interaction on R_global_, *F* = 0.59, *p* = .440.

### Diagnostic and Amyloid-status Group Differences in Direct vs. Indirect Connections

To gain a more thorough understanding of how diagnosis and A*β*-status affect hippocampal and LFC redundancy measures, the constituent properties of redundancy, including direct (i.e., R_direct_) and indirect connections (i.e., R_indirect_), were separately analyzed.

### Direct Connections

First considering R_direct_ of the posterior hippocampi, there were no diagnostic differences in R_direct_ of the left posterior HC, *F* = 0.51, *p* = .479, or the right posterior HC, *F* = 0.001, *p* = .972. There were trend-level A*β*-status differences in R_direct_ in the left posterior HC, *F* = 3.34, *p* = .068, and significant A*β* differences in the right posterior HC, *F* = 5.59, *p* = .018. In both the left and right posterior HC, A*β*- exhibited higher R_direct_ than A*β*+. There was a trend-level interaction of diagnosis and A*β*-status on R_direct_ for the left posterior HC, *F* = 3.23, *p* = .075. Post hoc analyses revealed that CN A*β*- exhibited higher direct connections with the left posterior HC than CN A*β*+, *p* = .013 (FDR corrected). The interaction between diagnosis and A*β*-status on R_direct_ for the right posterior HC was not significant, *F* = 0.05, *p* = .820.

For the anterior hippocampi, there were significant diagnostic differences in R_direct_ for both the left anterior HC, *F* = 4.66, *p* = .034, and right anterior HC, *F* = 5.13, *p* = .025. In both regions, MCI exhibited more direct connections with the bilateral anterior HC than CN. There were neither A*β*-status differences (*F* = 0.27, *p* = .606 for left anterior HC, *F* = 0.76, *p* = .382 for right anterior HC) nor interactions of diagnosis and A*β*-status in R_direct_ for either left or right anterior HC (*F* = 1.32, *p* = .250 for left anterior HC, *F* = 0.73, *p* = .396 for right anterior HC).

For the LFC, there were no diagnostic differences, *F* = 0.05, *p* = .823, no A*β*-status differences, *F* = 1.29, *p* = .252, and no interactions of diagnosis and A*β*-status in R_direct_, *F* = 0.13, *p* = .727.

### Indirect Connections

For R_indirect_, similar effects as R_direct_ were observed in each region of interest. Beginning with the posterior hippocampi, there were no diagnostic differences in R_indirect_ of the left posterior HC, *F* = 0.20, *p* = .651, or the right posterior HC, *F* = 0.003, *p* = .955. However, there were A*β*-status differences in R_indirect_ for the left posterior HC, *F* = 4.64, *p* = .032, and the right posterior HC, *F* = 6.15, *p* = .012, such that A*β*- had a higher number of indirect connections with both regions than A*β*+. There were no interactions between diagnosis and A*β*-status on R_indirect_ in the left posterior HC, *F* = 2.25, *p* = .134, or right posterior HC, *F <* 0.001, *p* = .995.

For the anterior hippocampi, there were trend-level diagnostic differences in R_indirect_for the left anterior HC, *F* = 3.61, *p* = .055, such that MCI had a higher number of indirect connections than CN. There were no diagnostic differences in R_indirect_ for the right anterior HC, *F* = 2.04, *p* = .158. There were neither A*β*-status differences (*F* = 0.13, *p* = .716 for left anterior HC, *F* = 1.22, *p* = .281 for right anterior HC) nor interactions of diagnosis and A*β*-status on R_indirect_ (*F* = 1.09, *p* = .306 for left anterior HC, *F* = 0.81, *p* = .371 for right anterior HC).

For the LFC, there were no diagnostic differences, *F* = 1.74, *p* = .183, no A*β*-status differences, *F* = 0.0003, *p* = .985, and no interactions of diagnosis and A*β*-status in R_direct_, *F* = 0.59, *p* = .443.

### Moderation by Diagnosis and Amyloid-status on SUVR-Redundancy Relationship

Though the A*β*- group showed greater R_global_ than A*β*+ across CN and MCI groups primarily in the posterior hippocampi, we further examined whether the relationships between the level of A*β* deposition, measured continuously through SUVR, and redundancy differed by A*β*-status within each diagnostic group or by diagnosis. There were significant moderation effects of the SUVR-R_global_ relationship in right anterior and left posterior HC regions. Results from the corresponding regression models can be found in Tables 2 and 3 and Figure 4. Regression parameters for other regions (i.e., left anterior HC, right posterior HC, and LFC) are reported in the Supplementary Material.

**Figure 4.**
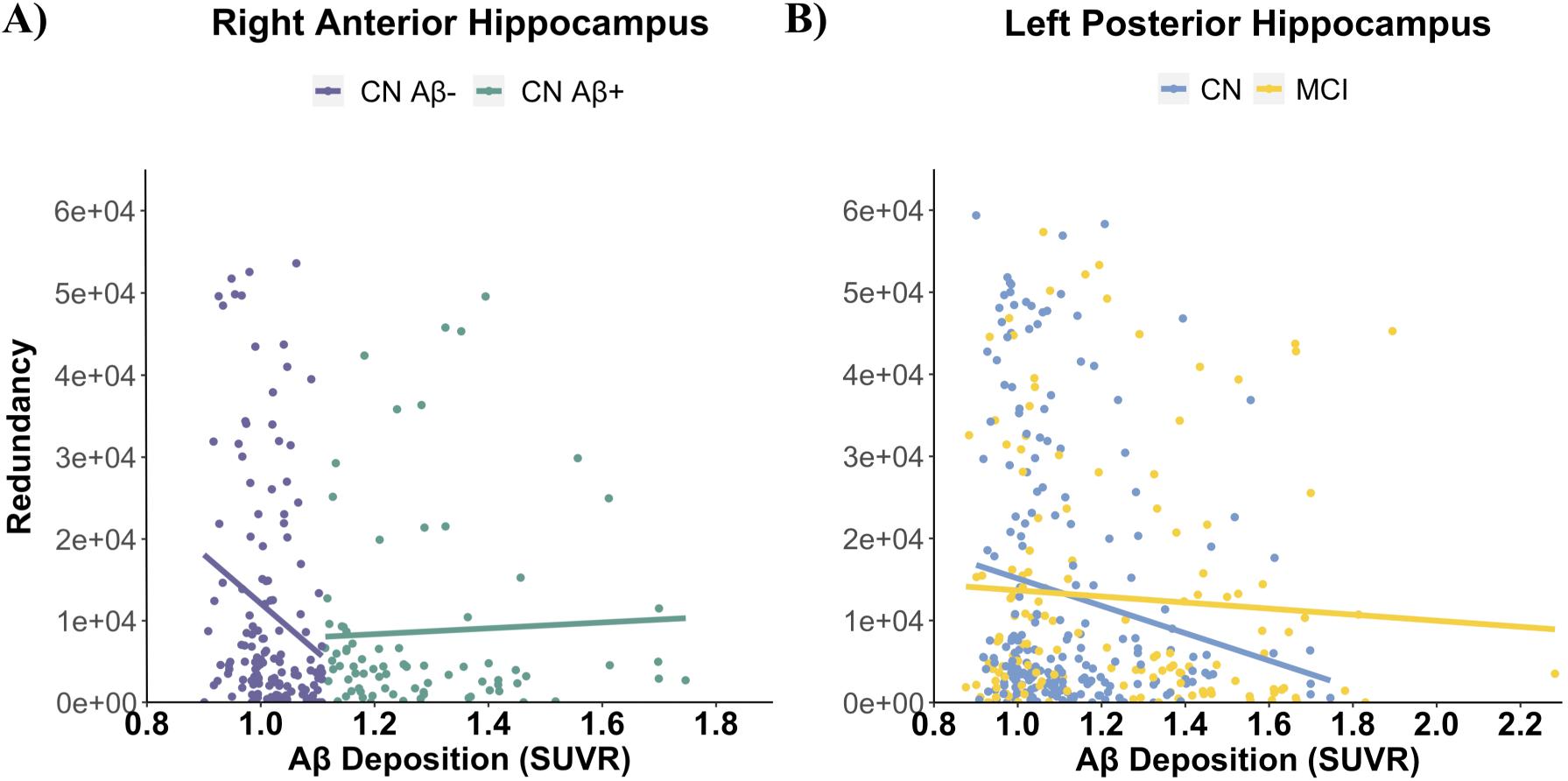
Moderators of the Aβ deposition and hippocampal redundancy relationship. (A) In CN, Aβ- exhibited a strong negative relationship between SUVR and right anterior hippocampal redundancy. This effect was not observed in Aβ+. (B) In the right posterior hippocampus, CN showed a strong negative relationship between SUVR and redundancy, which was not observed in MCI.

**Table 2.** Moderation regression parameters for the right anterior hippocampus.

| Group | R<sup>2</sup> / Model | Predictors | $\beta$ | Uncorrected | Corrected |
| --- | --- | --- | --- | --- | --- |
|  | *p*-value |  |  | *p*-value | *p*-value |
| CN | R<sup>2</sup> = 0.09<br>*p* = .005 | Age | 0.188 | 0.012* | 0.062<sup>+</sup> |
|  |  | Sex | -0.35 | 0.011* | 0.062<sup>+</sup> |
|  |  | Education | -0.04 | 0.565 | 0.682 |
|  |  | APOE4 | 0.029 | 0.846 | 0.935 |
|  |  | SUVR | -0.97 | 0.005** | 0.062<sup>+</sup> |
| | | Aβ-status | 0.386 | 0.159 | 0.364 |
| | | SUVR*Aβ-status | 0.951 | 0.012* | 0.062<sup>+</sup> |
| MCI | R<sup>2</sup> = 0.026<br>*p* = .828 | Age | -0.003 | 0.974 | 0.974 |
|  |  | Sex | 0.014 | 0.94 | 0.974 |
|  |  | Education | -0.067 | 0.444 | 0.621 |
|  |  | APOE4 | -0.216 | 0.309 | 0.468 |
|  |  | SUVR | 0.659 | 0.227 | 0.397 |
| | | Aβ-status | -0.473 | 0.312 | 0.468 |
| | | SUVR*Aβ-status | -0.748 | 0.185 | 0.364 |
|  | R<sup>2</sup> = 0.024<br>*p* = .271 | Age | 0.09 | 0.114 | 0.341 |
|  |  | Sex | -0.189 | 0.09<sup>+</sup> | 0.313 |
|  |  | Education | -0.029 | 0.585 | 0.682 |
|  |  | APOE4 | -0.068 | 0.576 | 0.682 |
|  |  | SUVR | -0.177 | 0.062<sup>+</sup> | 0.26 |
|  |  | Diagnosis | 0.168 | 0.135 | 0.355 |
|  |  | SUVR*Diagnosis | 0.149 | 0.191 | 0.364 |

Note. <sup>+</sup>*p* < .10; \**p* < .05; \*\**p* < .01.

**Table 3.**
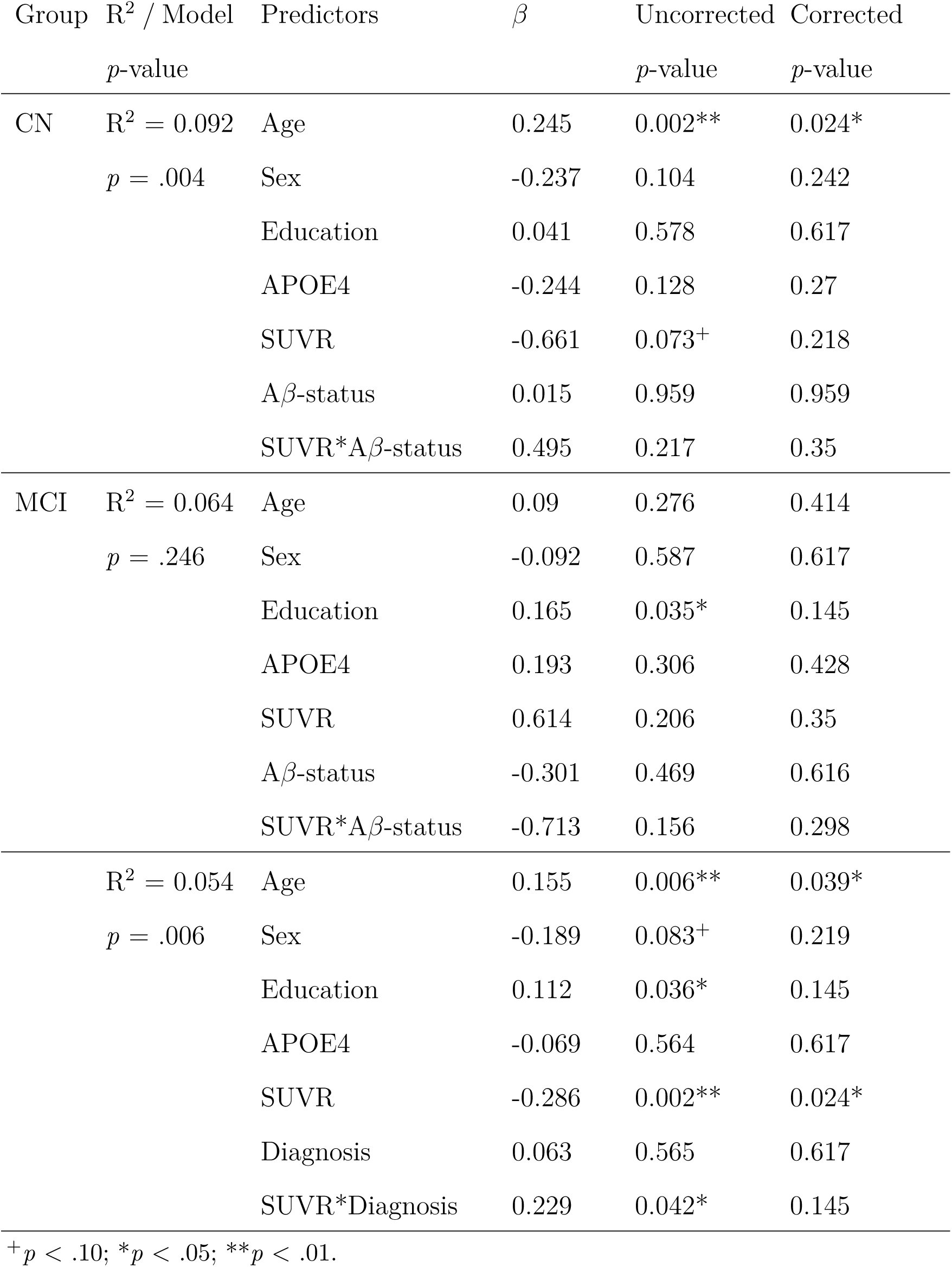
Moderation regression parameters for the left posterior hippocampus.

#### Amyloid-status Moderates Right Anterior HC Amyloid SUVR-Redundancy Relationships in Cognitively Normal Older Adults

In CN, A*β*-status was found to moderate the relationship between A*β* level and right anterior HC R_global_, *β* = 0.951, *p* = .0012 (Figure 4A; Table 2). Posthoc investigation revealed that higher A*β* SUVR was associated with lower right anterior HC R_global_ in A*β*- (*p* = .01), while this relationship was not significant for A*β*+ (*p* = .90). The relationship between A*β* level and redundancy was not moderated by A*β*-status in the left anterior HC, *β* = 0.303, p = .417, left posterior HC, *β* = 0.495, p = .217, right posterior HC, *β* = 0.526, p = .187, or LFC, *β* = -0.378, p = .297. In MCI, A*β*-status did not significantly moderate the relationship between A*β* level and R_global_ in any regions, *p*’s *>* .05 (Tables 2 and 3).

#### Diagnosis Moderates Left Posterior HC Amyloid SUVR-Redundancy Relationships

Diagnosis was found to moderate the relationship between A*β* level and left posterior HC R_global_, *β* = 0.229, *p* = .042 (Figure 4B; Table 3). Simple slopes analysis revealed that higher A*β* SUVR was associated with lower left posterior HC R_global_ in CN (*p <* .001), while this relationship was not significant for MCI (*p* = .41). Diagnosis did not moderate the relationship between A*β* level and R_global_ in the left anterior HC, *β* = 0.138, *p* = .229, right anterior HC redundancy, *β* = 0.149, *p* = .191, right posterior HC, *β* = 0.115, *p* = .303, or LFC redundancy, *β* = 0.132, *p* = .246.

### Relationship between Redundancy and Years of Education

To examine the potential role of redundancy as a resilience factor, the relationships between R_global_, in each ROI, and years of education, a proxy of cognitive resilience, were tested. Analyses were collapsed across diagnostic groups. There was a significant positive relationship between years of education and left posterior HC redundancy (*β* = 0.12, *p* = .022) and a trend-level positive relationship between years of education and right posterior HC redundancy (*β* = 0.098, *p* = .062) (Figure 5A,B). Higher R_global_ of the left and right posterior HC was associated with higher years of education. Redundancy in the left anterior HC, *β* = -0.031, *p* = .557, right anterior HC, *β* = -0.030, *p* = .564, and LFC, *β* = -0.064, *p* = .221, were not related to years of education.

**Figure 5.**
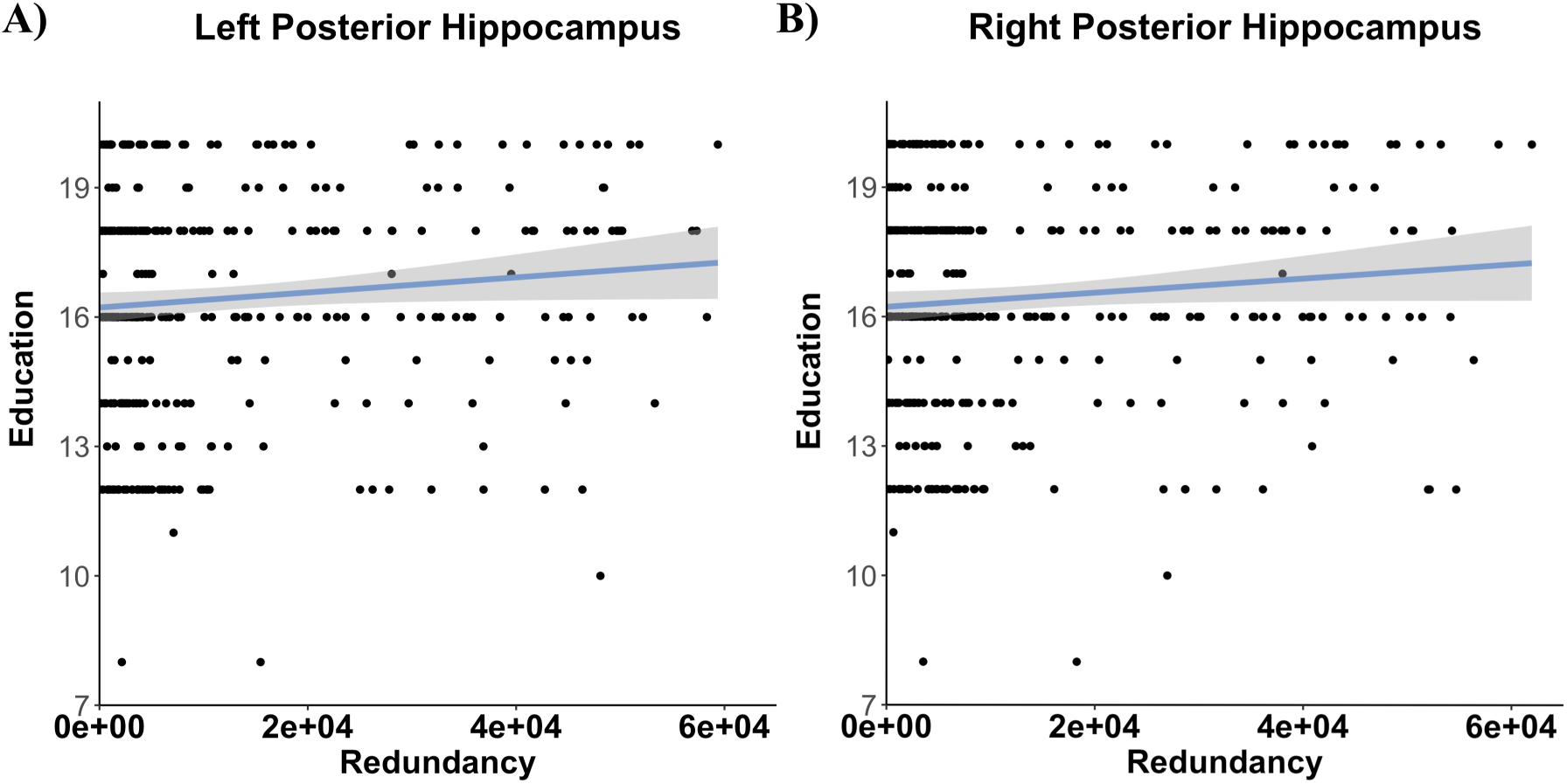
Posterior hippocampal redundancy and years of education. Years of education was positively related to (A) left posterior hippocampal redundancy, and (B) right posterior hippocampal redundancy.

### Posterior HC Redundancy Related to Better Memory Performance

To identify if redundancy confers a cognitive-advantage, redundancy-cognition relationships were explored. Replicating past work (Langella et al., 2021), we found that R_global_ in the left posterior HC, *β* = 0.134, *p* = .007, and right posterior HC, *β* = 0.167, *p* = .001, were related to memory performance, such that higher R_global_ was associated with better memory performance across CN and MCI groups (Figure 6A,B). To further probe this relationship, we conducted regressions evaluating the relationship between memory and R_global_ of the left and right posterior HC separately in A*β*- and A*β*+ groups. We found that the positive relationship between memory and left posterior HC redundancy was evident in A*β*-, *β* = 0.183, *p* = .004, but not A*β*+, *β* = -0.002, *p* = .982 (bottom right Figure 6A). Similar trends were found for the right posterior HC, such that the positive relationship between memory and right posterior HC redundancy was evident in A*β*-, *β* = 0.163, *p* = .011, but not A*β*+, *β* = 0.109, *p* = .177 (bottom right Figure 6B). There was no significant relationship between memory performance and R_global_ in the left anterior HC, *β* = -0.080, *p* = .102, right anterior HC, *β* = -0.043, *p* = .388, or LFC, *β* = -0.045, *p* = .366. Further, no relationships were observed between executive function performance and R_global_ in any ROI (*β* = -0.012, *p* = .819 for left posterior HC *β* = 0.013; *p* = .802 for right posterior HC; *β* = -0.049, *p* = .332 for left anterior HC; *β* = -0.032, *p* = .523 for right anterior HC; *β* = -0.051, *p* = .312 for LFC).

**Figure 6.**
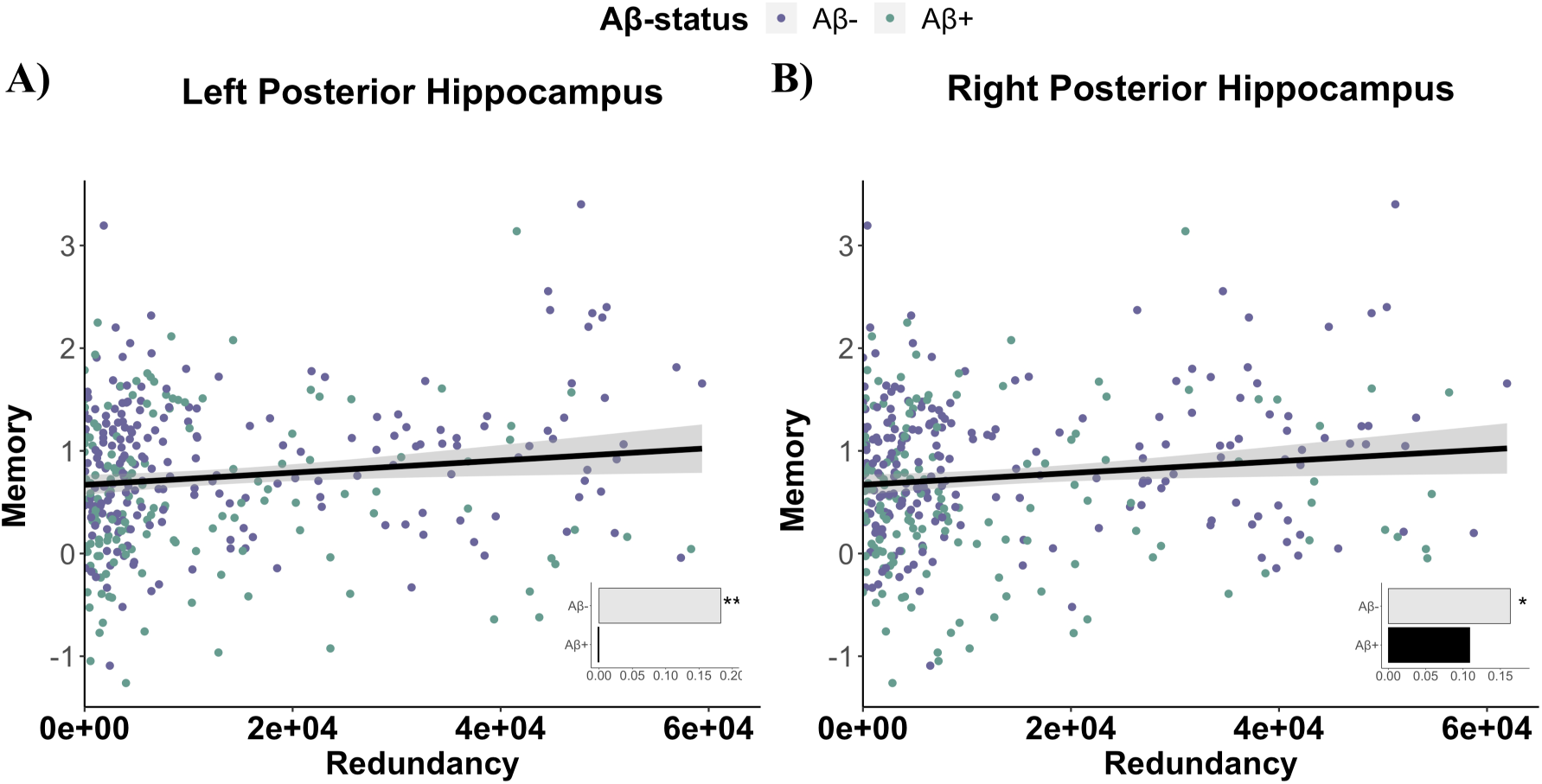
Posterior hippocampal redundancy and memory performance. Higher memory performance was associated with higher (A) left posterior hippocampal redundancy, and (B) right posterior hippocampal redundancy.

### Redundancy Moderates Pathology-Cognition Relationships

Finally, we explored whether redundancy measures moderate the relationship between pathology and cognition. Analyses were conducted separately for CN and MCI groups and for memory and executive composite scores. For memory composite scores, the CN group did not show any moderation effect of redundancy on the relationship between A*β*-SUVR and memory performance in any ROI (*β* = 0.029 *p* = .698 for left posterior HC; *β* = 0.041, *p* = .580 for right posterior HC; *β* = 0.019, *p* = .769 for left anterior HC; *β* = 0.052 *p* = .422 for right anterior HC; *β* = -0.011, *p* = .881 for LFC). In MCI, left posterior HC redundancy was found to moderate the SUVR-memory relationship, at trend, *β* = -0.13, *p* = .085, such that high left posterior HC redundancy (+1SD) was associated with a stronger negative relationship between A*β* deposition and memory performance (*p <* .001; Figure 7A). Redundancy of the right posterior HC, *β* = -0.075, *p* = .411, left anterior HC, *β* = 0.011, *p* = .891, right anterior HC, *β* = 0.017, *p* = .820, and LFC, *β* = 0.006, *p* = .942, did not moderate the relationship between A*β* level and memory performance in MCI.

**Figure 7.**
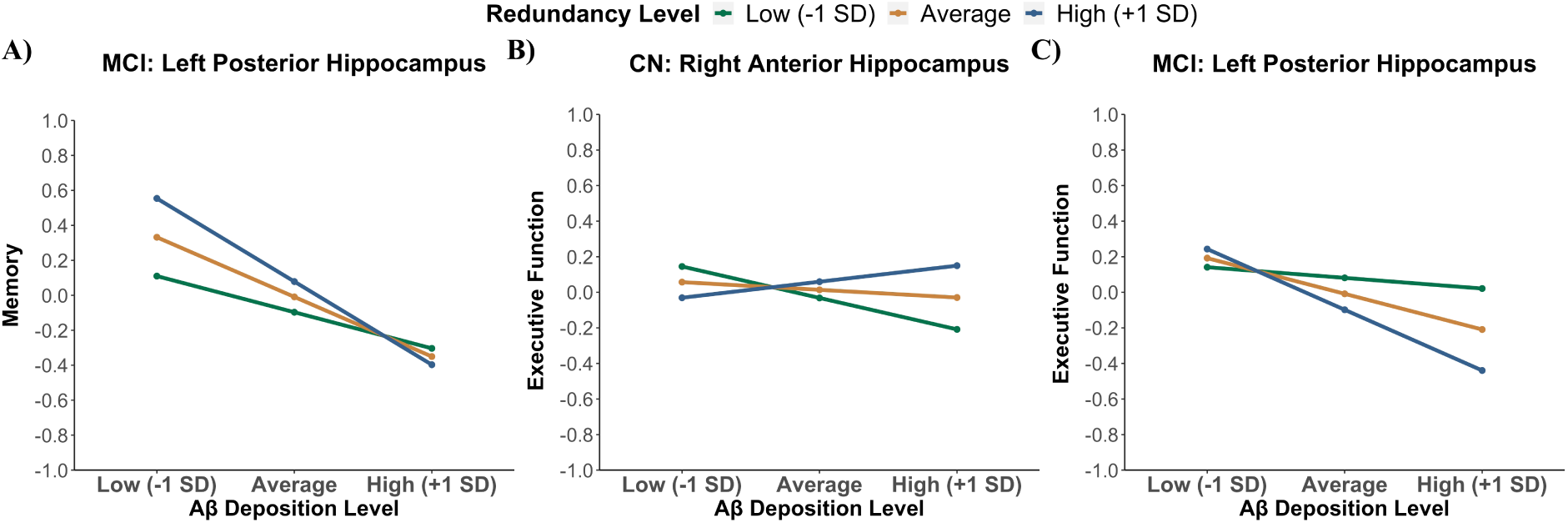
Moderators of Aβ deposition and cognition relationships. (A) In MCI, left posterior hippocampal redundancy moderated the relationship between Aβ deposition and memory performance, at trend. (B) In CN, right anterior hippocampal redundancy moderated the relationship between Aβ deposition and executive function performance. (C) In MCI, left posterior hippocampal redundancy moderated the relationship between Aβ deposition and executive function performance, at trend.

For executive function performance, the CN group showed a trend-level moderation effect of right anterior HC redundancy on the SUVR-executive function relationship *β* = 0.134, *p* = .052. Low right anterior HC redundancy (−1SD) was associated with a significant negative relationship between A*β* deposition and memory performance (*p <* .001), but this effect was not observed with HC redundancy at the mean or at high levels (+1SD) (Figure 7B). Redundancy of the left anterior HC, *β* = -0.009, *p* = .897, left posterior HC, *β* = 0.014, *p* = .867, right posterior HC, *β* = -0.007, *p* = .930, and LFC, *β* = 0.030, *p* = .699, did not moderate the relationship between A*β* deposition and executive function performance in CN. In MCI, there was a trend-level moderation effect of left posterior HC redundancy on A*β* SUVR-executive function relationship, *β* = -0.141, *p* = .080 (Figure 7C). High left posterior HC (+1SD) was associated with a significant negative relationship between A*β* deposition and executive function performance (*p*= .01), a relationship which was not observed for low left posterior HC redundancy (*p*= .60). Redundancy of the left anterior HC, *β* = -0.076, *p* = .350, right anterior HC, *β* = -0.082, *p* = .282, right posterior HC, *β* = -0.140, *p* = .138, and LFC, *β* = -0.066, *p* = .415, did not moderate the relationship between A*β* level and executive function performance in MCI.

## Discussion

Understanding the mechanisms underlying resilience in brain aging have significant implications for early intervention and prevention strategies in AD. In the current investigation, we examined whether regional functional redundancy confers resilience against developing AD neuropathology. Using graph theory, redundancy was quantified in five *a priori* defined ROIs previously implicated in brain aging resilience (i.e., left and right anterior HC, left and right posterior HC, LFC) by summarizing the direct (path length = 1) and indirect (path length = 2-4) paths connecting each set of regions in the brain. Our results revealed several noteworthy findings: (1) Higher A*β*-level was associated with lower posterior hippocampal redundancy, independent of diagnosis; (2) the relationship between A*β* status and posterior HC redundancy was similar for both direct and indirect connections; (3) a diagnosis of MCI was related to higher anterior hippocampal redundancy; (4) higher posterior HC redundancy related to better memory performance and higher years of education, a proxy of cognitive resilience, especially among A*β*- individuals; (5) right anterior HC redundancy may act as a cognitive resilience factor, particularly in preserving executive function performance in the earliest stages of A*β* accumulation, and (6) functional redundancy does not underlie the resilience effects of global LFC connectivity.

Previous studies have shown a disruption of functional redundancy in MCI patients compared to cognitively normal older adults (Langella et al., 2021). It remained unresolved whether functional redundancy was affected by A*β* levels in the preclinical and prodromal stages of AD. We observed that A*β*-positivity selectively disrupted redundancy of the left and right posterior HC, but not anterior HC or LFC. The A*β*-related effects were evident across direct and indirect portions of redundancy for bilateral posterior HC, though A*β* accumulation had a stronger effect on indirect than direct connections for the left posterior HC. The redundancy metric is a comprehensive metric of regional connectivity, in that it captures not only the integrity of the connectivity profile of the ROI itself (e.g., left posterior HC direct connections), but also the integrity of the connectivity profiles of the graphical neighbors of the ROI (e.g., precuneus, retrosplenial cortex). By separately assessing direct and indirect connections of each ROI, we showed that A*β* disrupts not only connectivity of the posterior hippocampus but also that of its direct neighbors.

It is intriguing that A*β*-related decreases in functional redundancy were regionally specific to the posterior HC, but not other ROIs, such as the anterior HC. Previous studies found that two cortical networks exist and converge within the hippocampus including the anterior temporal (AT) and posterior medial (PM) systems (Ranganath & Ritchey, 2012). The anterior HC is strongly connected to regions of the AT system, such as the anteriolateral entorhinal cortex, perirhinal cortex, amygdala, ventral temperopolar cortex, and lateral orbitofrontal cortex and contributes to processing object/item based components of episodic memory (Kahn, Andrews-Hanna, Vincent, Snyder, & Buckner, 2008; Libby, Ekstrom, Ragland, & Ranganath, 2012; Ranganath & Ritchey, 2012). On the other hand, the posterior HC is strongly connected to PM regions, including the posteriomedial entorhinal cortex, parahippocampal cortex, retrosplenial cortex, posterior cingulate cortex (PCC), and precuneus and underscores processing contextual information of episodic memory (Kahn et al., 2008; Libby et al., 2012; Ranganath & Ritchey, 2012). The AT and PM systems are further differentiated by their selective susceptibility to AD pathology, as regions of the AT network are most burdened by tau pathology while the PM network is most vulnerable to A*β* pathology (Dautricourt et al., 2021; Maass et al., 2019). The susceptibility of the PM system to A*β* pathology is supported by the extant AD literature, which demonstrates that major hubs of the PM, including the precuneus and posterior cingulate cortex, that show a high degree of connectivity throughout the brain (van den Heuvel & Sporns, 2013), are some of the earliest regions to exhibit and drive A*β* pathology deposition (Buckner, Andrews-Hanna, & Schacter, 2008; Palmqvist et al., 2017; Pasquini et al., 2017). A major consequence of such regional A*β* accumulation is cascading network failure (Jones et al., 2016) that ultimately disrupts connectivity between these posterior hubs, the posterior HC (Gardini et al., 2015; Jones et al., 2016; Qi et al., 2010; Tahmasian et al., 2015), and throughout the brain (Jones et al., 2016). Collectively, these findings suggest that regional redundancy is a rich single-scalar metric that captures brain-wide disruptions to functional connectivity, which occur as AD neuropathology develops, particularly within the posterior hippocampal network. Given the susceptibility of the anterior hippocampal network to tau pathology, it will be important in future investigations to explore redundancy-tau relations across the AD continuum.

Replicating past work (Langella et al., 2021), we found that posterior, but not anterior, HC conferred a cognitive benefit, such that higher left or right posterior HC redundancy were related to better episodic memory performance. We extend these findings by demonstrating that this relationship was primarily driven by A*β*- groups. Specifically, we found a significant positive relationship between bilateral HC redundancy and memory performance in A*β*- but this relationship was not significant in A*β*+. These findings may suggest that the presence of abnormal brain A*β* in posterior networks decouples the functional cognitive benefit conferred by posterior HC redundancy from memory performance, regardless of clinical status.

In regard to the anterior HC, we observed that redundancy levels were not affected by A*β*-status. This finding is in-line with the reported distinct vulnerability of anterior and posterior hippocampal systems to AD pathology (Dautricourt et al., 2021; Maass et al., 2019), such that the posterior networks are selectively targeted by A*β* pathology while anterior networks are vulnerable to tau pathology (Adams, Maass, Harrison, Baker, & Jagust, 2019; Maass et al., 2019). Though there were no amyloid effects on anterior HC redundancy, we did find evidence of diagnostic differences, particularly when the global redundancy measure was segregated into direct and indirect components. Specifically, we observed that MCI had, on average, higher levels of direct connections between the left or right anterior HC and all other brain regions compared to CN. Consistent with these findings, a recent study showed that MCI and AD patients exhibited hyperconnectivity between the anterior hippocampus and the AT system, which may be driven by tau accumulation (Berron et al., 2021; Dautricourt et al., 2021). Without measures of tau pathology, it is difficult to discern the affiliated or underlying neurobiological alterations that may drive increased anterior hippocampal direct connections in MCI. However, it is likely that the detrimental effects of A*β* on posterior systems paired with advancing MTL tau pathology mediate system-level network reorganization in MCI. Together, these findings suggest that changes to anterior hippocampal networks are rather robust to A*β* but evidence from the literature suggests that such systems are selectively vulnerable to tau-related changes that may occur following A*β* accumulation in the posterior system.

Compared to posterior HC redundancy, we found that redundancy of the LFC did not differ by diagnosis or A*β*-status. This finding is analogous to past work examining functional connectivity of the LFC (Franzmeier et al., 2017, 2018), and suggests that LFC connectivity is relatively unaffected by developing amyloid pathology or progression across the AD continuum. The resilience of the LFC network to AD pathological changes makes it well-positioned to serve as a neural mechanism of resilience to facilitate the recruitment of additional neural networks to potentially compensate for network failure that occurs as a result of accumulating pathology. Our results of no association between LFC redundancy and diagnosis or A*β*-status suggest that the resilience mechanism supported by LFC connectivity could differ from those supported by functional redundancy.

To determine if regional redundancy exhibits properties of a resilience factor, we examined the relationship between regional redundancy and years of education that is commonly used as a proxy of cognitive resilience. We found that only redundancy within the left and right posterior HC, but not anterior HC or LFC, were positively related to years of education. These findings provide insight that posterior HC redundancy may be shaped by early life experiences and that it has the potential to play a role in resilience of brain aging (Langella et al., 2021). To further determine if regional redundancy confers cognitive resilience to developing AD neuropathology, we tested if redundancy moderated the relationship between A*β* deposition and cognitive performance (i.e., memory, executive function). Our findings did not support the hypothesis that the left or right posterior HC redundancy mitigated the potential deleterious effect of A*β* pathology on cognition. Contrary to our prediction, in MCI, we observed that the negative influence of A*β* on memory and executive function was exacerbated by high levels of left posterior HC redundancy, though this effect was marginal. This finding is in line with recent work which demonstrates that hyperconnectivity of the posterior hippocampus and parahippocampal cortex with medial PFC in MCI is associated with worse cognitive performance (Gardini et al., 2015). Together, these results may indicate that neural hyperconnectivity, which occurs in the posterior hippocampal network during the MCI stage, may represent aberrant neural activity that reflects the deleterious effect of pathology, rather than the compensatory role, on cognition.

Although at trend level, we found that the negative relationship between A*β* load and executive function performance was mitigated in those with high levels of right anterior HC redundancy among cognitively normal older adults. The adverse effect of A*β* on executive function is well-documented (Baker et al., 2017; Tideman et al., 2021) and has been shown to emerge in the preclinical phase, when A*β* accumulation is dominating the neocortex. Due to the significant connections between the anterior hippocampus and frontal lobes (Ranganath & Ritchey, 2012), one explanation is that the maintained functionality within this network, particularly in the preclinical phase (i.e., normal cognition but high levels of cortical A*β*), confers protection for frontally-supported cognition, such as executive function. However, the results were significant at trend level, thus need to be interpreted with caution.

In addition to the posterior HC, we found that redundancy of the LFC did not moderate the relationship between A*β* pathology and cognition. Past work has shown that the global connectivity profile of the LFC confers cognitive resilience in pathological aging (Franzmeier et al., 2017, 2018). We examined whether LFC redundancy (i.e., repetitive functional wiring between LFC and all other brain regions) may serve a similar protective role as the global connectivity measure of the LFC. Our redundancy measures of LFC, however, did not replicate the previous findings of the LFC’s role in conferring cognitive resilience in the presence of A*β* pathology. These results together suggest that it may not be the total number of connections, either direct or indirect, with the LFC but rather the strength of the direct connections with LFC that underlie its role in resilience.

Several limitations to the current study need to be noted. First, our ability to draw conclusions concerning the temporal relations between A*β* deposition and disrupted posterior hippocampal redundancy and relative changes across the AD continuum were limited by the cross-sectional design. For example, we interpret that reduced posterior HC redundancy may occur as a result of preferential deposition and the subsequent disruption of posterior networks by A*β*. However, alternative explanations cannot be ruled out due to the cross-sectional design. Rather, it could be that lower hippocampal redundancy marks inefficient neural processing. Coupled with increasing age and other biological risk factors (e.g., *APOE4*), such inefficiency may tax the system and drive vulnerability for A*β* deposition. It will be important in follow up studies to utilize a longitudinal design to illuminate the temporal relationship between posterior hippocampal redundancy and vulnerability to A*β*. To expand on promising previous work illustrating the potential role of posterior HC redundancy as a cognitive resilience factor in aging (Langella et al., 2021), we considered A*β* as our primary marker of AD pathology. However, given the particular sensitivity of the anterior HC network to tau accumulation (Maass et al., 2019) and the synergistic effects of A*β* and tau to disrupt functional networks (Berron et al., 2021), it will be important in future studies to examine both pathological hallmarks in the context of resilience. An additional limitation includes the limited diversity of the sample. It is well-documented that AD disproportionately affects under-represented populations, in particular Black Americans (Gurland et al., 1999; Rajan, Weuve, Barnes, Wilson, & Evans, 2019; Steenland, Goldstein, Levey, & Wharton, 2016). Mounting evidence suggests that early life factors, such as years or quality of education, significantly contributes to racial disparities in cognitive levels, including episodic memory (Sisco et al., 2015; Yaffe et al., 2013). Considering that posterior hippocampal redundancy, in particular, is related to memory performance and may be shaped by early life experiences, as evidenced by its positive association with years of education, it will be critically important to examine such a metric in an ethnically-diverse sample. These limitations, however, should not underscore the strengths of the current study, including a well-powered sample size and further characterization of the role of regional redundancy in pathological brain aging.

In summary, regional redundancy represents a rich metric that captures not only the functional integrity of the primary region of interest but also the functional integrity of the region’s graphical neighbors. Due to this rigorous dimensionality, redundancy measures can capture global functional network disruptions that occur as a consequence of AD pathology, and may show potential as a marker of pathological brain aging.

## Acknowledgments

Data collection and sharing for this project was funded by the Alzheimer’s Disease Neuroimaging Initiative (ADNI) (National Institutes of Health Grant U01 AG024904) and DOD ADNI (Department of Defense award number W81XWH-12-2-0012). ADNI is funded by the National Institute on Aging, the National Institute of Biomedical Imaging and Bioengineering, and through generous contributions from the following: AbbVie, Alzheimer’s Association; Alzheimer’s Drug Discovery Foundation; Araclon Biotech; BioClinica, Inc.; Biogen; Bristol-Myers Squibb Company; CereSpir, Inc.; Cogstate; Eisai Inc.; Elan Pharmaceuticals, Inc.; Eli Lilly and Company; EuroImmun; F. Hoffmann-La Roche Ltd and its affiliated company Genentech, Inc.; Fujirebio; GE Healthcare; IXICO Ltd.; Janssen Alzheimer Immunotherapy Research & Development, LLC.; Johnson & Johnson Pharmaceutical Research & Development LLC.; Lumosity; Lundbeck; Merck & Co., Inc.; Meso Scale Diagnostics, LLC.; NeuroRx Research; Neurotrack Technologies; Novartis Pharmaceuticals Corporation; Pfizer Inc.; Piramal Imaging; Servier; Takeda Pharmaceutical Company; and Transition Therapeutics. The Canadian Institutes of Health Research is providing funds to support ADNI clinical sites in Canada. Private sector contributions are facilitated by the Foundation for the National Institutes of Health (www.fnih.org). The grantee organization is the Northern California Institute for Research and Education, and the study is coordinated by the Alzheimer’s Therapeutic Research Institute at the University of Southern California. ADNI data are disseminated by the Laboratory for Neuro Imaging at the University of Southern California. This reserach was additionally supported by NIH R01AG068990 (H. Oh), NIH R01AG069265 (W. Heindel/S. Buka), and NIH S10OD025181 (J. Sanes).

## Notes

https://adni.loni.usc.edu/

### Competing Interest Statement

The authors have declared no competing interest.

