## Supplemental Tables for "Functional Redundancy of the Posterior Hippocampi, but not Anterior Hippocampi or Left Frontal Cortex, is Disrupted in Pathological Brain Aging"

### Supplementary Material

**Supplementary Table 1**. Left anterior hippocampus.

| Group | $R^2$ / Model $p$-value | Predictors | $\beta$ | Uncorrected $p$-value | Corrected $p$-value |
|---|---|---|---|---|---|
| CN | $R^2 = 0.034$ | Age | 0.127 | $0.085^{+}$ | 0.553 |
| | $p = .380$ | Sex | -0.223 | $0.099^{+}$ | 0.553 |
| | | Education | -0.015 | 0.829 | 0.958 |
| | | APOE4 | 0.057 | 0.7 | 0.918 |
| | | SUVR | -0.369 | 0.282 | 0.845 |
| | | A$\beta$-status | 0.088 | 0.746 | 0.922 |
| | | SUVR*A$\beta$-status | 0.303 | 0.417 | 0.917 |
| MCI | $R^2 = 0.026$ | Age | -0.075 | 0.437 | 0.917 |
| | $p = .818$ | Sex | 0.32 | 0.105 | 0.553 |
| | | Education | -0.081 | 0.369 | 0.917 |
| | | APOE4 | -0.037 | 0.867 | 0.958 |
| | | SUVR | 0.353 | 0.533 | 0.918 |
| | | A$\beta$-status | -0.193 | 0.691 | 0.918 |
| | | SUVR*A$\beta$-status | -0.363 | 0.534 | 0.918 |
| | $R^2 = 0.015$ | Age | 0.028 | 0.627 | 0.918 |
| | $p = .620$ | Sex | 0.003 | 0.979 | 0.979 |
| | | Education | -0.023 | 0.673 | 0.918 |
| | | APOE4 | 0.004 | 0.977 | 0.979 |
| | | SUVR | -0.114 | 0.229 | 0.803 |
| | | Diagnosis | 0.204 | $0.07^{+}$ | 0.553 |
| | | SUVR*Diagnosis | 0.138 | 0.229 | 0.803 |

Note. Regression parameters for the moderation of the A$\beta$ deposition and left anterior hippocampus relationship by A$\beta$-status and DX. $^{+}p < .10$; $^{*}p < .05$; $^{**}p < .01$.

**Supplementary Table 2**. Right posterior hippocampus.

| Group | $R^2$ / Model *p*-value | Predictors | $\beta$ | Uncorrected *p*-value | Corrected *p*-value |
|---|---|---|---|---|---|
| CN | $R^2 = 0.091$ | Age | 0.266 | 0.001** | 0.017* |
| | $p = .005$ | Sex | -0.044 | 0.761 | 0.841 |
| | | Education | -0.008 | 0.918 | 0.918 |
| | | APOE4 | -0.335 | 0.036* | 0.129 |
| | | SUVR | -0.766 | 0.037* | 0.129 |
| | | A$\beta$-status | 0.236 | 0.417 | 0.614 |
| | | SUVR*A$\beta$-status | 0.526 | 0.186 | 0.49 |
| MCI | $R^2 = 0.083$ | Age | 0.093 | 0.26 | 0.546 |
| | $p = .101$ | Sex | 0.104 | 0.539 | 0.666 |
| | | Education | 0.167 | 0.032* | 0.129 |
| | | APOE4 | 0.149 | 0.431 | 0.614 |
| | | SUVR | 0.375 | 0.439 | 0.614 |
| | | A$\beta$-status | -0.293 | 0.481 | 0.632 |
| | | SUVR*A$\beta$-status | -0.495 | 0.324 | 0.567 |
| | $R^2 = 0.058$ | Age | 0.17 | 0.002** | 0.025* |
| | $p = .003$ | Sex | 0.013 | 0.905 | 0.918 |
| | | Education | 0.085 | 0.109 | 0.327 |
| | | APOE4 | -0.143 | 0.235 | 0.546 |
| | | SUVR | -0.253 | 0.007** | 0.046* |
| | | Diagnosis | 0.052 | 0.637 | 0.743 |
| | | SUVR*Diagnosis | 0.115 | 0.303 | 0.567 |

Note. Regression parameters for the moderation of the A$\beta$ deposition and right posterior hippocampus relationship by A$\beta$-status and DX. $^{+}p < .10$; $*p < .05$; $**p < .01$.

**Supplementary Table 3**. Left frontal cortex.

| Group | $R^2$ / Model *p*-value | Predictors | $\beta$ | Uncorrected *p*-value | Corrected *p*-value |
|---|---|---|---|---|---|
| CN | $R^2 = 0.044$ | Age | 0.164 | 0.023* | 0.478 |
| | $p = .215$ | Sex | -0.007 | 0.958 | 0.992 |
| | | Education | -0.071 | 0.294 | 0.497 |
| | | APOE4 | 0.145 | 0.315 | 0.497 |
| | | SUVR | 0.226 | 0.497 | 0.58 |
| | | A$\beta$-status | -0.126 | 0.633 | 0.7 |
| | | SUVR*A$\beta$-status | -0.378 | 0.297 | 0.497 |
| MCI | $R^2 = 0.037$ | Age | 0.086 | 0.379 | 0.497 |
| | $p = .635$ | Sex | 0.266 | 0.187 | 0.497 |
| | | Education | -0.083 | 0.37 | 0.497 |
| | | APOE4 | -0.21 | 0.351 | 0.497 |
| | | SUVR | 0.756 | 0.192 | 0.497 |
| | | A$\beta$-status | -0.603 | 0.225 | 0.497 |
| | | SUVR*A$\beta$-status | -0.717 | 0.232 | 0.497 |
| | $R^2 = 0.028$ | Age | 0.106 | $0.061^{+}$ | 0.497 |
| | $p = .178$ | Sex | 0.106 | 0.338 | 0.497 |
| | | Education | -0.058 | 0.285 | 0.497 |
| | | APOE4 | -0.001 | 0.992 | 0.992 |
| | | SUVR | -0.076 | 0.419 | 0.517 |
| | | Diagnosis | 0.133 | 0.235 | 0.497 |
| | | SUVR*Diagnosis | 0.132 | 0.246 | 0.497 |

Note. Regression parameters for the moderation of the A$\beta$ deposition and left frontal cortex relationship by A$\beta$-status and DX. $^{+}p < .10$; $^{*}p < .05$; $^{**}p < .01$.
